# Selective modulation of the glucocorticoid receptor with CORT108297 during chronic adolescent stress evokes sex-specific effects in adulthood

**DOI:** 10.1101/646745

**Authors:** Evelin Cotella, Rachel Morano, Aynara Wulsin, Susan Martelle, Paige Lemen, Maureen Fitzgerald, Benjamin A. Packard, Rachel Moloney, James Herman

## Abstract

Adolescent animals are vulnerable to the effects of stress on brain development. We hypothesized that long-term effects of adolescent chronic stress are mediated by glucocorticoid receptor (GR) signaling. We used a specific GR modulator (CORT108297) to pharmacologically disrupt GR signaling in adolescent rats during exposure chronic variable stress (CVS). Male and female rats received 30mg/kg of drug concomitantly with a 2-week CVS protocol starting at PND46. Emotional reactivity (open field) and coping behaviors (forced swim test (FST)) were then tested in adulthood, 5 weeks after the end of the CVS protocol. Blood samples were collected two days before FST and serial samples after the onset of the swim test to determine baseline and stress response levels of HPA hormones respectively.

Our results support differential behavioral, physiological and stress circuit reactivity to adolescent chronic stress exposure in males vs. females, with variable involvement of GR signaling. In response to adolescent stress, males had heightened reactivity to novelty and exhibited marked reduction in neuronal excitation following swim stress in adulthood, whereas females developed a passive coping strategy and enhanced HPA axis stress reactivity. Only the latter effect was attenuated by treatment with the GR modulator C108297. Our data suggest that adolescent stress differentially affects emotional behavior and circuit development in females, and that GR plays a role in driving some but not all sequelae of adolescent stress.

## Introduction

Traumatic experiences during development are associated with diverse psychopathologies such as depression and anxiety disorders as well as hyphtolamic-pituitary adrenal (HPA) axis dysregulation (Oitzl et al., 2010). The adolescent brain is considered particularly susceptible to changes in environmental conditions (Brenhouse and Andersen, 2011). Areas involved in cognitive and affective processes, such as hippocampus, prefrontal cortex (PFC) and amygdala, are still developing their adult structure and function during this period by undergoing processes of synaptic pruning, myelination, and changes in the expression of key regulatory molecules (Andersen and Teicher, 2008, 2004; Brenhouse and Andersen, 2011). Several authors have proposed this phase as a critical period for the development of vulnerability to psychiatric disorders later in life (Eiland and Romeo, 2013; Leussis and Andersen, 2008). Moreover, traumatic or chronic stress during this phase is associated with the onset of psychiatric disorders (Goodyer, 2000; Patel et al., 2007).

The actions of glucocorticoids in the brain and other organs are modulated by signaling through corticosteroid receptors, GR (glucocorticoid receptor) and MR (mineralocorticoid receptor) (Reul and de Kloet, 1985). These receptors contribute to regulate the HPA axis stress response by negative feedback mechanisms (de Kloet et al., 2008). In rodents, early life stress as well as chronic exposure to stress in adulthood lastly decrease the expression of GR (Cattaneo and Riva, 2016; de Kloet et al., 2005; Herman et al., 2005), which has been linked to reduced inhibition of HPA axis stress responses. Of note, chronic exposure to glucocorticoids, acting through GR signaling, is linked to dendritic retraction, reduced cell growth and (in hippocampus) reduced neurogenesis (reviewed in (Joëls et al., 2007)), and it is thought that signaling through GRs plays a role triggering lasting effects of early life experience (reviewed in (Champagne et al., 2009; Herbert et al., 2006)). Notably, GR is expressed abundantly in the hippocampus, amygdala, and prefrontal cortex during adolescent development (as well as adulthood) (Eiland and Romeo, 2013; Herman et al., 2005; Herman and Cullinan, 1997), indicating that these structures may be susceptible to fluctuations in glucocorticoids during adolescent stress exposure.

Prior data from our group indicates that chronic variable stress (CVS) during adolescence can evoke changes in anxiety like-behavior and coping style in both males and females. In the case of the HPA axis response to stress, females showed reduced stress responsivity and males were not affected (Cotella et al., 2019; Wulsin et al., 2016). To further understand the mechanisms behind these sex-specific enduring effects of adolescent stress, we tested the hypothesis that adolescent chronic stress causes life-long, sex-specific effects that are determined by the signaling through the glucocorticoid receptor (GR) during stress exposure. We used a recently-developed GR specific modulator (CORT108297) to target GR signaling during the exposure to adolescent stress, in order to test the involvement of GR in defining the long-term effects previously observed in our model.

## Materials and Methods

Male and female Sprague Dawley rats were bred in-house, weaned at postnatal day 21 (PND21) and pair-housed in standard clear cages (20 cm height ×22cm width × 43cm length)) under a 12 h light/ 12 h dark cycle (lights on at 7:00 am) at constant room temperature (23±2 °C) with *ad libitum* access to food and water. No more than 2 littermates were included in each experimental group. All tests were performed during the light cycle, between 09:00 AM and 2:00 PM. All procedures and care performed in the animals were approved by the University of Cincinnati Institutional Animal Care and Use Committee.

### Adolescent chronic variable stress

Rats in the CVS group were subjected to 14 days of our standard CVS paradigm during late adolescence (PND 46 ± 2) following previous work from our group (Jankord et al., 2011; Wulsin et al., 2016) with some modifications. The CVS protocol included a set of unpredictable variable stressors applied twice daily (AM and PM): 1) 1h shaker stress (100rpm), 2) 1h cold room (4°C), 3) 30 minute restraint, 4) 30 min hypoxia exposure (8% O2 and 92% N2) 5) 1h wet bedding. Swimming was not used as a CVS stressor to ensure that FST was a novel stressor. Following CVS, animals were allowed to recover for 5 weeks to be evaluated during adulthood. We did not include overnight stressors to avoid extending the time of exposure to stressors too far from the injection time. The timeline of the experiment is shown in Figure 1.A.

**Figure 1:**
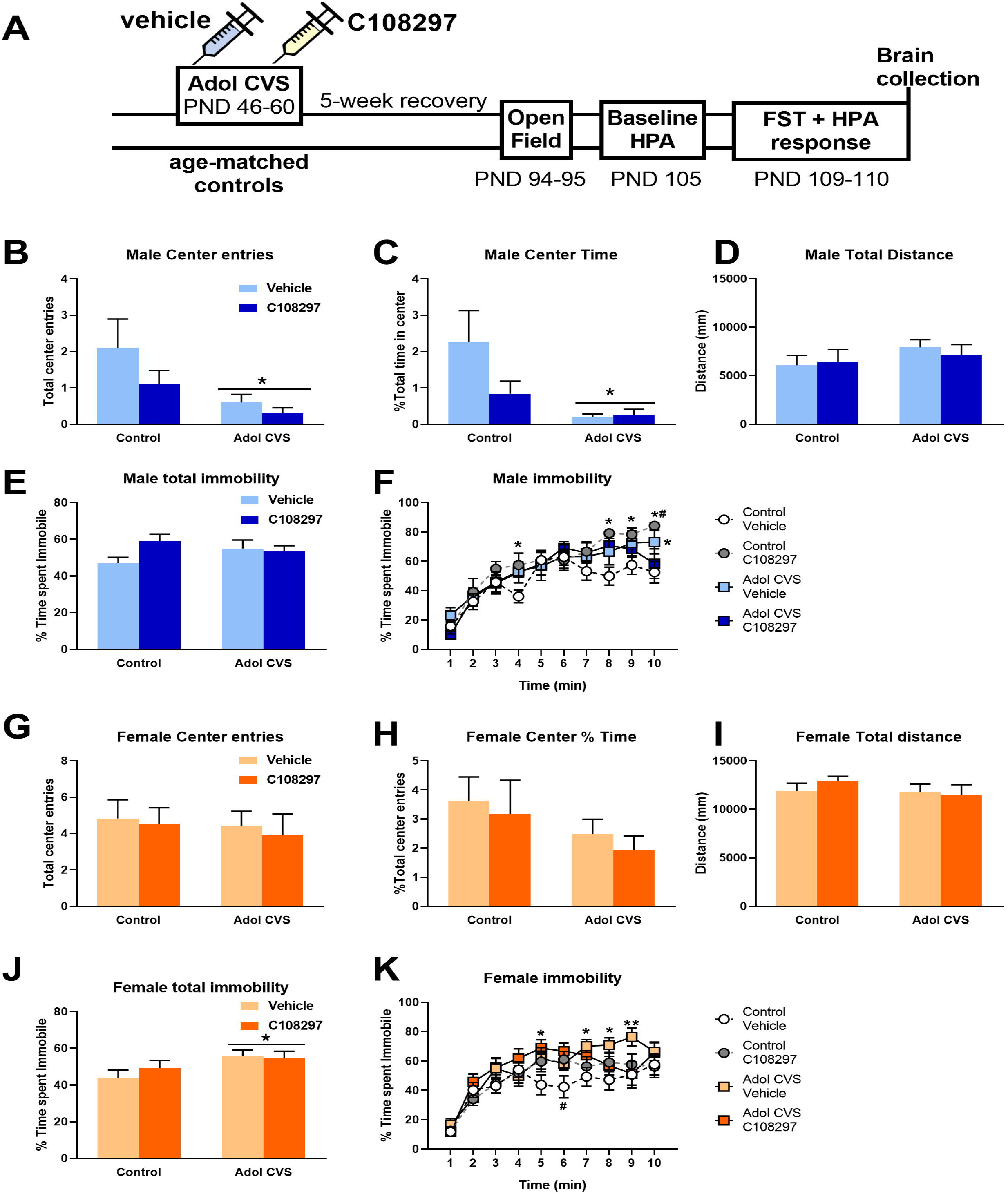
A: Experimental timeline Animals were subjected to adolescent CVS (adol) and concomitantly administered with CORT108297 (30mg/Kg) or vehicle during 2 weeks starting at PND 46, and evaluation started after 5 weeks of recovery. **Activity in the open field test arena after 5 min exploration**: Total number of entries (**B**: males; **G**: females), Total time in the center (**C**: males; **H**: females), total distance covered (**D**: males; **I**: females). Data are presented as mean ± s.e.m. *: significant result p < 0.05. **Total percent of immobility in the forced swim test (FST) after 10 min exposure** (**E**: males; **J**: females) and **percent of immobility over time** (**F**: males; **K**: females). Data are presented as mean ± s.e.m. **E, J**: *: significant result p < 0.05 versus corresponding control group. **F, K:** *: significant result p < 0.05 versus control vehicle group. ^#^: significant result p < 0.05 versus CVS C108297 group.

### GR specific modulation

CORT108297 (30mg/kg) (Corcept therapeutics) or vehicle (polyethylene glycol) was subcutaneously injected daily at 9:00 throughout the duration of the CVS protocol. AM stressors started between 30min to 1 h after the injection and PM stressors happened between 5 - 5.5 h after the injection (According to reports from Corcepts, the half-life of the compound is compatible with once-a-day oral dosing (CORCEPT THERAPEUTICS INC, 2011)). The resulting groups for each sex were: Control-Vehicle; Control-CORT108297; CVS-Vehicle and CVS-CORT108297.

### Somatic markers

Body weight was recorded daily during the duration if the CVS protocol to administer the proper dose of C108297 every day. After the conclusion of CVS, rats were weighed once a week to keep track of the general health status of the animals as well as for possible extended effect of the experimental conditions. Data are presented as total body weight, percent of body weight gain from the first day of experiment during CVS and percent of body weight gain from the last day of CVS, measured once a week, to assess the recovery after CVS. At euthanasia, internal organs that are known to reflect chronic stress effects (adrenals, thymus and heart) were collected and weighed.

### Open field evaluation

On PND94-95 rats were evaluated during a 5 minute exposure to an open field. Briefly, animals were placed on the corner of a 1m2 arena. Videos were acquired using Clever Capture Star software (CleverSys, Reston, VA) and then they were automatically scored using Clever Top Scan software (CleverSys, Reston, VA). The variables measured were total distance covered in the arena, total number of center entries and time spent in the center.

### Forced swim test

At days 109-110 rats were evaluated in the forced swim test during 10 min (25+2 °C). Videos were acquired with Clever Capture Star software (CleverSys, Reston, VA and manually scored by a blinded observer. The method used was sampling the behavior each 5s. Behaviors considered: climbing, swimming, immobility and diving.

### Blood sampling and hormone assessment

At PND 105 rats were quickly tail bled (<3min) to get samples to determine baseline levels of stress hormones. The day of the FST (PND109) rats were bled from their tails 15, 30, 60 and 120 minutes after the onset of the test to assess the HPA stress response to a novel stressor. Samples were obtained by quickly clipping the distal tip of the tail of a freely moving rat with a razor blade and collecting ∼250 ul of blood into EDTA (10ul, 100mM) containing tubes. Subsequent samples were obtained by gently removing the clot from the distal tail to recommence bleeding. All samples were collected within 3 min. In order to minimize hormonal variability due to circadian fluctuations, all procedures were performed during circadian nadir of the diurnal CORT rhythm and blood samples were collected before 1:00 PM. After collection, blood was centrifuged at 3500 x *g*, for 15 min at 4 °C and then plasma samples were extracted and kept at −20 °C until hormones level determination. Corticosterone (CORT) and adrenocorticotropic hormone (ACTH) plasma levels were measured by radioimmunoassay as previously described (Smith et al., 2018; Wulsin et al., 2016). CORT concentration was determined using 125I RIA kits (MP Biomedicals Inc., Orangeburg NY). ACTH was determined by radioimmunoassay, using 125I ACTH as trace (Amersham Biosciences) (ACTH antiserum donated by Dr. William Engeland (University of Minnesota) at a 1:120,000 dilution (Ulrich-Lai et al., 2006). The ratio of plasma corticosterone to the log of plasma ACTH was calculated as an index of adrenal sensitivity to ACTH (ENGELAND et al., 1981; Ulrich-Lai and Engeland, 2002).

### Immunohistochemistry

Immediately after the last serial blood sample was taken, rats were euthanized with an overdose of sodium pentobarbital and subsequently transcardially perfused with 0.9% saline followed 4% paraformaldehyde in 0.1 M phosphate buffer (PBS), pH 7.4. Brains were post-fixed in 4% paraformaldehyde at 4°C for 24h, then transferred to 30% sucrose in 0.01MPBS at 4°C. Brains were sliced into serial 35μm coronal sections using a freezing microtome (−20°C). Sections (1/12) were collected into wells containing cryoprotectant solution (30% Sucrose, 1% Polyvinyl-pyrolidone (PVP-40), and 30% Ethylene glycol, in 0.1 M PBS). For immunolabeling, sections were washed 6×5min in 0.01M PBS at room temperature (RT). After being rinsed, sections were incubated with 1% Sodium Borohydride in 0.1MPBS for 30 min at RT. After rinsing 6×5 min 0.1M PBS, they were incubated in 3% hydrogen peroxide diluted in 0.1M PBS for 20 min. Subsequently, brain slices were rinsed 6×5 min and 4×15 min in 0.1 M PBS and then incubated in blocking solution (4% normal goat serum (NGS), 0.4% TritonX-100, 0.2% bovine serum albumin (BSA) in 0.1M PBS, 2h at RT. Sections were then incubated with c-Fos rabbit polyclonal antibody (1:1000, Santa Cruz, sc-52) in blocking solution, overnight at RT. The next day, sections were rinsed (3 × 5 min) in 0.1M PBS at RT, followed by incubation with secondary antibody (biotinylated goat anti-rabbit, (1:400; Vector Laboratories, BA1000) in blocking solution at RT for 1h. Sections were again rinsed (3×5 min) in 0.1M PBS and then reacted with avidin-biotin horseradish peroxidase complex (1:800 in 0.1M PBS; Vector Laboratories) for 1 h at RT. Sections were then rinsed (3×5 min) in 0.1M PBS and then developed with a 8 min incubation in DAB-Nickel solution: 10mg 3,3′-diaminobenzidine (DAB) tablet (Sigma, DF905), 0.5 ml of a 2% aqueous nickel sulfate solution, 20ul of 30% hydorgen peroxide in 50ml of 0.1M PBS. Sections were finally washed in PBS, mounted on superfrost slides (Fisherbrand, Fisher), allowed to dry, dehydrated with xylene, and then coverslipped in DPX mounting medium (Sigma). Sections from 7 to 8 brains per experimental group were processed. For analysis, we counted 3 bilateral sections from equivalent coordinates covering the anterior, medial and posterior portions of the prefrontal cortex (PFC), nuclei in the amygdala: central amygdaloid nucleus (CeAm), medial amygdaloid nucleus (MeAm) and basolateral amygdala (BLA) and the subdivisions of the paraventricular nucleus of the thalamus (PVT): anterior (PVA), intermediate (PV) and posterior (PVP). In the case of the paraventricular nucleus of the hypothalamus (PVN) and the lateral ventral septum (LSV), we analyzed 4 anteroposterior bilateral sections from equivalent coordinates. Each brain region limit and coordinates were defined following a brain atlas (Paxinos and Watson, 2007). The number of Fos positive neuclei was counted with a semiautomatized method using ImageJ software (National Institutes of Health, Bethesda, MD). Counts of Fos immunoreactive cells were obtained from each area of interest using the Analyze Particle tool, using a defined common level of background intensity, nuclei circularity and size (previously validated manually). Once the number of Fos positive nuclei was determined in each section, the relative density of the population of immunopositive cells was calculated by dividing this number by the area measured in each case.

### Statistical analysis

Data were analyzed using STATISTICA 7.0 (Statsoft, Inc.,Tulsa, USA), Prism 8 (GraphPad Software, La Jolla California USA) and Infostat (Di Rienzo et al., n.d.). For the analysis of the results from OFT, FST, baseline levels of hormones, a 2-way ANOVA (2×2 design: STRESS x TREATMENT) analysis was performed with a level of significance p < 0.05. HPA axis stress response and behavior on the FST over time were analyzed by repeated measurements ANOVA, STRESS × TREATMENT × TIME, with a level of significance of p < 0.05. In the cases where significant differences were found, a Tukey post-hoc test was performed. In some cases, we performed planned comparison to evaluate individual differences in cases where there was not a significant interaction between factors. When necessary, data were processed by logarithmic or square root transformations to allow parametric analysis. In the case of open field, the results were analyzed using general linearized models (GLM). Only statistics for significant results are discussed in the results sections.

## Results

### Behavior

#### Open Field

The behavioral response to 5-minute exposure to an open field arena was analyzed as a measure of emotional reactivity. In males, prior adolescent CVS had a main effect reducing the total time spent in the center F_(1, 35)_= 6.10, p<0.05, with no effect of treatment nor interaction between factors. The planed comparisons showed that both CVS groups were significantly different from the control-vehicle group (p<0.05). The control-C108297 group was not different from the corresponding vehicle group, nor from the corresponding CVS group. Adol CVS also reduced the number of total entries to the center of the arena, F_(1, 35)_ = 6.44, p<0.05. There was no drug effect nor interaction between factors and no individual differences when analyzing planned comparisons. There was no effect of stress or drug on the time spent in the center zone in females, nor the total number of entries in it. Importantly, none of the factors had an effect on locomotor activity in both sexes. Effects shown in Figs. 1.B-D and 1.G-I.

### Forced swim test

Behavioral adaptation during the forced swim test is summarized in Figures 1.E-F and 1.J-K. There was no effect of stress, drug treatment or their interaction on total immobility during the test. However, responses are analyzed over time, and we observed a significant STRESSxTREATMENT interaction, F_(1, 36)_= 4.338, p<0.05. Although the interaction over time was not significant, F_(9, 324)_= 1.772, p=0.0726, we used planned comparisons to determine possible differential effects between groups. Increased immobility was observed in the CVS-vehicle group relative to the control-vehicle group only in the last time point (p<0.05). Meanwhile, the CVS-C108297 group was not different from the control-vehicle levels, suggesting a mild effect of the drug in preventing the effect of CVS. The control-C108297 group increased immobility relative to the control-vehicle group over several time points (p<0.05) and was significant with respect to the CVS-C108297 group at the last time point (p<0.05). The data are consistent with a minor impact of modulating GR signaling during adolescence on the behavioral response to an aversive situation during adulthood in males.

In females, there was an effect of adol CVS on total immobility F_(1, 44)_= 5.331, p<0.05 in the FST. A STRESS×TREATMENT×TIME interaction F_(9, 396)_= 2.357, p<0.05 was observed upon temporal analysis of FST behavior. In this analysis the CVS-vehicle group significantly differed from the control-vehicle group across several time points (p<0.05) near the end of the test, indicating a shift in the coping strategy of these animals compared to the corresponding controls. Furthermore, the CVS-vehicle group differed from the CVS-C108297 group only at one later time point (min=9, p<0.05), again suggesting a mild effect of GR modulation during adolescence on the effects of stress in adulthood in the females. The control animals treated with the drug during adolescence, similar to what was observed in the males, showed a significant increase of the immobility during minute 6 compared to the control vehicle group (p<0.05) but then they returned to similar levels as those of the corresponding control group.

### HPA axis

#### Baseline HPA activity

2 days before performing the forced swim test, a blood sample was obtained in order to determine the baseline levels of ACTH and corticosterone in the adult animals subjected to adolescent CVS and C108297 and their corresponding controls. When analyzing the effect on males, there was a significant interaction STRESSxTREATMENT F_(1, 32)_= 8.76, p<0.01 on baseline ACTH levels. The *post hoc* analysis revealed only a significant reduction of ACTH in the CVS-vehicle group compared to the corresponding unstressed group (p<0.05) (Fig 2.A). In the case of baseline corticosterone, C108297 had a main effect reducing the plasma levels of corticosterone F_(1, 36)_=6.88, p<0.05, with no effect of the stress protocol. The analysis of planned comparisons confirmed that both groups treated with C108297 had lower corticosterone levels than the corresponding vehicle groups (Fig 2.B).

**Figure 2:**
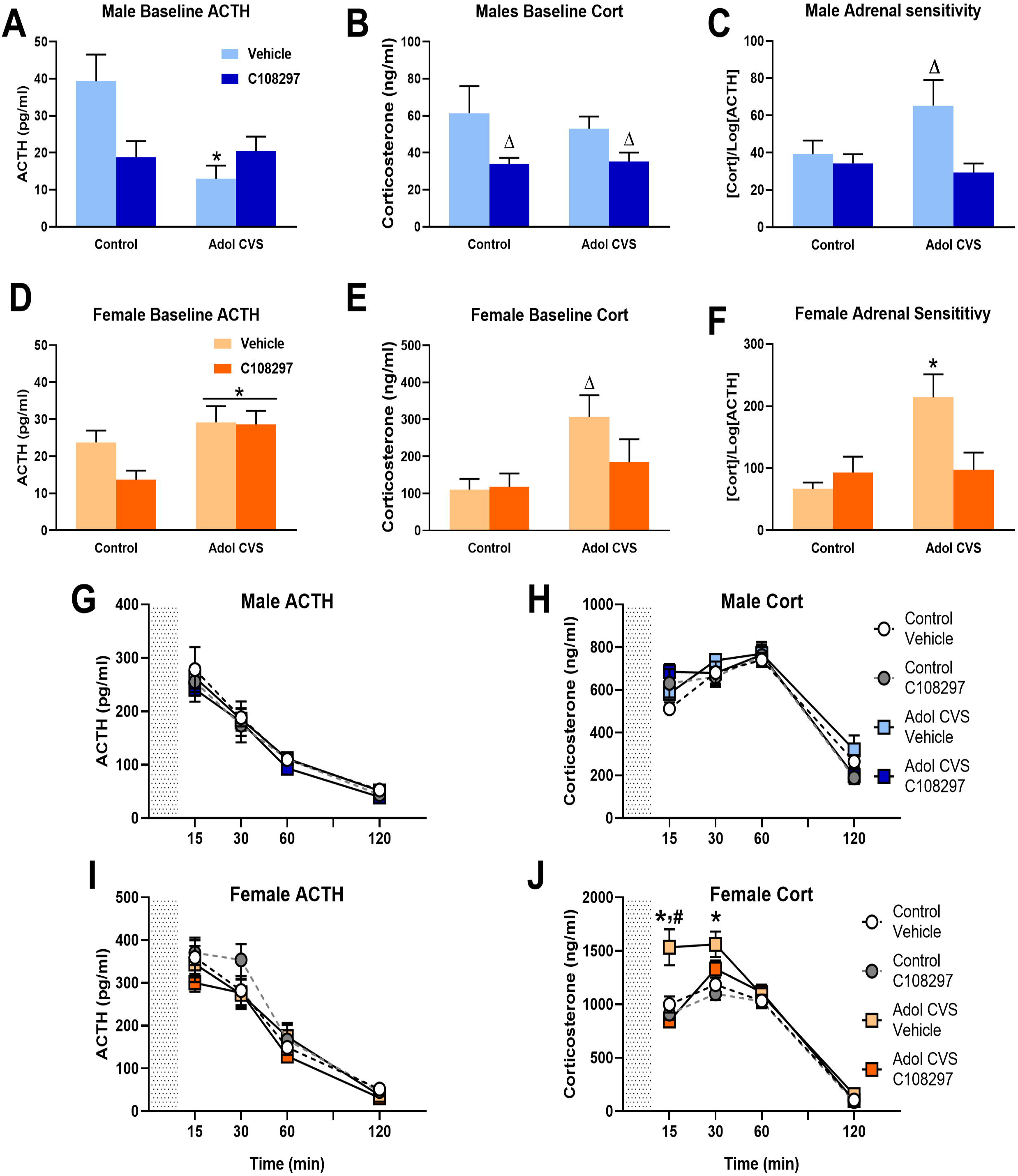
HPA axis regulation. Animals were subjected to adolescent CVS (adol) and concomitantly administered with CORT108297 (30mg/Kg) or vehicle during 2 weeks starting at PND 46, and evaluated after 6 weeks. **(A-F) Baseline hormones concentration and index of adrenal sensitivity in males and females**. Index of adrenal sensitivity was calculated as [Cort]/ log[ACTH] using baseline values Data are presented as mean ± s.e.m. *: significant result p < 0.05 versus corresponding unstressed. Δ: significant result p < 0.05 versus corresponding control or vehicle group (planned comparisons). **(G-J) Hormones concentration after 10 min exposure to FST** (grey area). Data are presented as mean ± s.e.m. *: significant result p < 0.05 versus corresponding control group. ^#^: significant result p < 0.05 vs CVS C108297.

In the case of the females, adol CVS had a main effect, increasing the levels of ACTH F_(1, 44)_= 7.23, p<0.05. In the case of corticosterone, there was a main effect of stress F_(1, 44)_=6.83, p<0.05. Planned comparisons to evaluate a possible effect of C108297 to prevent this effect showed that the CVS-vehicle group had significantly higher levels of corticosterone than its corresponding control-vehicle group (p<0.05) which was attenuated in CVS-C108297 group, which did not differ from control-vehicle of control-C108297 groups (Figs. 2.D or 2.E).

#### Index of adrenal sensitivity

The index of adrenal sensitivity, calculated as the concentration of baseline corticosterone over the logarithm of the baseline levels of ACTH. This gives an idea of how responsive the adrenal cortex is to the effects of ACTH (Engeland et al., 1981). In males, there was a main effect TREATMENT, F_(1,31)_= 6.88, p<0.05. This determined that in general, adolescent treatment with C108297 reduced adrenal sensitivity. Planned comparisons revealed significantly higher adrenal sensitivity in the CVS-vehicle group than the rest of the groups (p<0.05). There were main effects of both STRESS and TREATMENT in females (F_(1,38)=_8.09, p< 0.01 and F_(1,38)_=7.55, p<0.01, respectively). The STRESSxTREATMENT interaction was also significant F_(1,38)_= 5.88, p<0.05. There was a significant increase of the index in the CVS-vehicle group compared to the control groups (p<0.05) (Tukey’s test), and this was effectively prevented by the treatment with C108297 during adolescence CVS (p<0.05). Results are shown in figures 2.C and 2.F.

#### HPA response to a novel acute stressor

In order to assess the HPA stress response in the adult animals subjected to adolescent stress and C108297 treatment and their corresponding controls, after being evaluated in the forced swim test, serial blood samples were taken at 15, 30, 60 and 120 minutes after the onset of the test. In males, here were no effects of stress or drug on ACTH and corticosterone in any of the experimental groups (Figs 2.G and 2.H). In the case of females, there were main effects of STRESS F_(1, 44)_=14.57, p<0.001 and TREATMENT F _(1, 44)_=11.63, p<0.01 on corticosterone response, as well as multiple interactions: STRESSxTIME F_(3, 132)_=3.839, p<0.015; TREATMENT×TIME F_(3, 132)_=7.212, p<0.001; STRESS×TREATMENT F_(1, 44)_=5.063, p<0.05; STRESS×TREATMENT×TIME F_(3, 132)_= 4.534, p<0.01. In contrast, there was no effect of stress or treatment on ACTH release. *Post hoc* analysis indicated that the adol CVS-vehicle group had an exacerbated secretion of corticosterone at the 15 and 30 minute time points compared to the corresponding control group (p<0.05). Furthermore, treatment with C108297 significantly reduced this response at the 15 minute time point, indicating that treatment with C108297 during adolescent CVS prevented dysregulation of the HPA axis in adulthood in females (Figs 2.I and 2.J).

### Somatic markers of stress

#### Body weight

Body weight changes during CVS, post-CVS and across the whole experiment are summarized in Fig. 3. In males, there were significant main effects of STRESS F_(1,36)_=66.004, p<0.001, TREATMENT F_(1.36)_=14.791, p<0.001 and TIME F_(12,432)_=1534.22, p<0.001 during CVS. There were also significant STRESSxTIME F_(12,432)_=10.87, p<0.001 and TREATMENTxTIME F_(12,432)_=10.04, p<0.001 interactions. Planned comparisons indicated that treatment with C108297 decreased body weight gain in both adol CVS and control rats from day 7 to the end of CVS, suggesting an important metabolic effect of the drug itself. The CVS-vehicle group also showed reduced gain, consistent with a lasting chronic stress effect, from day 4 to the end of CVS. To accurately account for effects on normal body weight gain, the CVS-C108297 group was compared to the control vehicle group, given that C108297 alone affects body weight. The CVS-C108297 group had reduced weight gain relative to controls everyday from day 3 to the end of CVS. Finally, the CVS-C108297 group showed reduced weight gain compared to the corresponding control-C108297 group from day 3 to the end of CVS, suggesting an additive effect of drug and stress on metabolism (p<0.05) (Fig 3.A).

**Figure 3:**
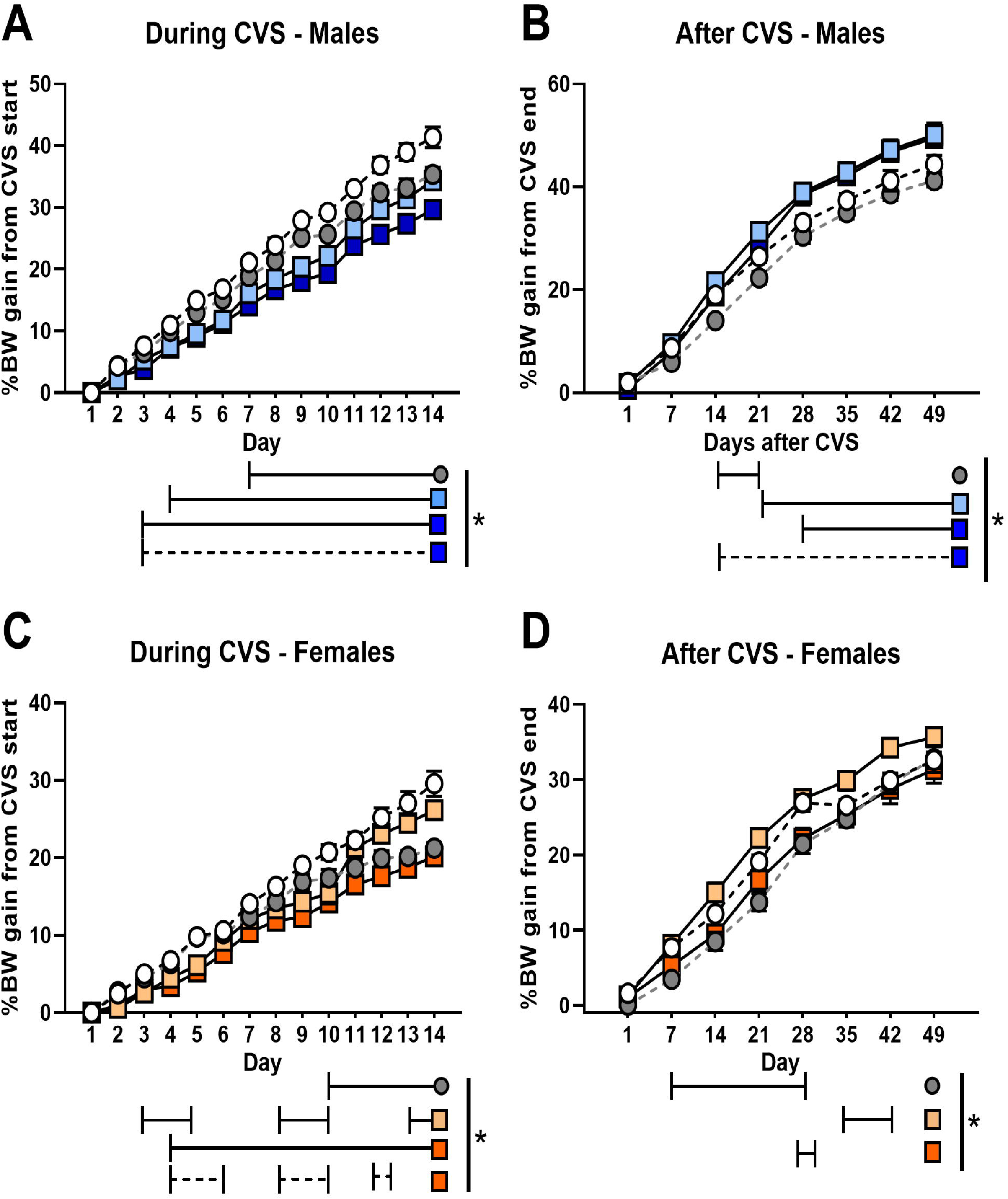
Body weight gain during (A, C) and after (B, D) CVS. Animals were subjected to adolescent CVS (adol) and concomitantly administered with CORT108297 (30mg/Kg) or vehicle during 2 weeks starting at PND 46. Body weight was taken weekly since before the beginning of CVS and daily during CVS (% weight gain calculated over CVS first day). After CVS, weight was taken weekly and % of weight gain was calculated from the last day of CVS. Data are presented as mean ± s.e.m. *: significant result p< 0.05 for planned comparisons or Tukey’s test (D). Solid lines represent comparisons versus Control-vehicle group. Dashed lines represent comparisons versus Control-C108297 group.

There was as significant main effect of STRESS on weight gain during recovery from CVS F_(1,36)_=20.01, p<0.001, indicating that both groups subjected to CVS increased rate of body weight gain after cessation of CVS. There was also an effect of TIME effect F_(7, 252)_=1581.106, p<0.001 and a significant STRESSxTIME interaction F_(7,252)_=12.26, p<0.001 in males. The control group treated with C108297 reduced weight gain rate only between 14 and 21 days after CVS compared to the control-vehicle group (p<0.05). On the contrary, the CVS-vehicle group and the CVS-C108297 group had continuous increased weight gain from day 21 and 14 respectively (p<0.05) comparing to the control-vehicle group. This is suggesting a metabolic compensatory effort in the animals subjected to CVS to recover growth rate (Fig. 3.B).

Despite the increase observed in the weight gain rate observed in the males after CVS, this was not reflected in absolute changes in body weight in the stressed group treated with C108297 since the groups treated with the compound had reduced absolute weight throughout the experiment (Supplemental Figure 1.A).

In females(Fig. 3.C), there were main effects of STRESS F_(1,44)_=18.005, p< 0.001, TREATMENT F_(1,44)_=16.638, p<0.001 and TIME F_(12,528)_=901.39, p<0.001 on body weight gain during CVS, due to a general reduction in weight gain seem both in the CVS and C108297 treated females (p<0.05). There were significant STRESS×TIME F_(12,528)_=4.248, p=0.001; TREATMENT×TIME F_(12,528)_ =24.208, p<0.001 interactions. Weight gain was reduced in both C108297 groups. The control-C108297 group had had reduced gain weight from day 10 until the end of the experiment (p<0.05). The CVS-vehicle group showed reduced weight gain between days 3 to 5, 8 to 10 and 13 to 14 (p<0.05) (suggesting possible cyclic effect related to interaction of stress with hormonal fluctuations in the females). Decreased weight gain was most pronounced in the CVS-C108297 group (reduced weight gain from day 4 on (p<0.05)), again suggesting an additive effects of stress and drug on metabolism. Weight gain in the CVS-C108297 group differed from the corresponding unstressed group on days 4 to 6, 8 to 10 and day 12 (p<0.05).

There were main effects of TREATMENT F_(1,44)_ =18.229, p< 0.001 and TIME F_(7,308)_=813.12, p<0.00000001 and weight gain after CVS, as well as several interactions: TREATMENT×TIME F_(7,308)_=3.818, p<0.001, TREATMENT×STRESS×TIME F_(7,308)_=2.316, p<0.05. *Post hoc* analysis indicate that controls treated with C108297 had reduced weight gain from days 7 until 21 after the end of CVS (p<0.05). Animals subjected to adol CVS and vehicle increased growth rate, but only for a limited time between days 35 and 42 (p<0.05). The CVS-C108297 group had reduced weight gain at day 28 after CVS (p<0.05), and did not differ from the control-C108297 group. Thus, the effect of the drug seems to have prevented the rebound on weight gain evoked by CVS (Fig. 3.D).

In the case of absolute body weight, C108297 evoked a reduction throughout the experiment (Supplemental Figure 1.B).

### Other somatic effects

Wet weights of the adrenal glands, thymus and heart was measured to account for effects of chronic stress. Since there was a difference in the body weight of the animals across the whole experiment due to the drug, the effect of adolescent CVS was analyzed by covariating the organs’ weight and the body weight measurement at the end of experiment.

The table shows the effects of stress and treatment on the somatic measurements. We did not observe effect of stress or C108297 on adrenal gland or thymus weight in males, nor covariation with the body weight. In the case of the heart, there was a main effect of STRESS F_(1,34)_=6.15, p<0.05 and positive covariation to body weight F_(1,34)_=18.82, p<0.001, correlation coefficient=1.99. Planned comparisons indicated that CVS evoked a long-term increase in the mass of the heart, regardless of the treatment. Results are shown in supplemental table 1.

**Table 1:** Fos immunoreactivity in the females. The table only shows the regions with negative results. Animals were subjected to adolescent CVS (adol) and concomitantly administered with CORT108297 (30mg/Kg) or vehicle during 2 weeks starting at PND 46. Brains were collected 2h after the onset of FST, after 6 weeks of recovery from adol CVS.

| Table 1 | Female Fos Immunoreactivity |  |  |  |
| --- | --- | --- | --- | --- |
| Brain region<br>(Fos-ir positive<br>nuclei/area) | Control-Vehicle | Control-C108297 | CVS-Vehicle | CVS-C108297 |
| PL (Fos-ir/mm<sup>2</sup>) | 133.32 ± 15.12 | 166.62 ± 30.25 | 122.78 ± 12.87 | 140.88 ± 26.61 |
| IL (Fos-ir/0.1 mm<sup>2</sup>) | 13.77 ± 1.8 | 14.41 ± 2.11 | 12.473 ± 1.54 | 14.17 ± 2.4 |
| PVN (Fos-ir/0.01 mm<sup>2</sup>) | 45.96 ± 8.80 | 43.20 ± 11.72 | 28.14 ± 7.73 | 33.16 ± 8.62 |
| MeAm (Fos-ir/0.1 mm<sup>2</sup>) | 17.01 ± 2.11 | 16.92 ± 2.98 | 13.8 ± 1.50 | 17.90 ± 2.16 |
| CeAm (Fos-ir/0.1 mm<sup>2</sup>) | 5.072 ± 1.09 | 5.98 ± 1.42 | 4.466 ± 0.81 | 4.49 ± 0.83 |
| BLA (Fos-ir/0.1 mm<sup>2</sup>) | 4.38 ± 0.82 | 6.59 ± 1.30 | 4.939 ± 0.58 | 5.50 ± 0.66 |
| PVA (Fos-ir/0.01 mm<sup>2</sup>) | 28.77 ± 5.99 | 26.29 ± 6.21 | 27.89 ± 3.72 | 38.85 ± 11.37 |
| PV (Fos-ir/0.01 mm<sup>2</sup>) | 34.56 ± 3.08 | 27.67 ± 5.69 | 24.37 ± 3.69 | 36.15 ± 6.58 |
| PVP (Fos-ir/0.01 mm<sup>2</sup>) | 30.41 ± 3.72 | 23.05 ± 5.15 | 32.18 ± 5.30 | 27.89 ± 6.43 |

In females, adrenal weights were not affected by STRESS or TREATMENT and were not affected by body weight differences. Regarding the thymus, there were no effects due to the experimental variables but there was a positive correlation with body weight F_(1,40)_=6.13, p<0.05, coefficient=1.29. Finally, regarding the heart, there were no experimental effects in the females, only positive correlation with body weight F_(1,42)_=26.76, p<0.001, coefficient=2.51. (Supplemental Table 1).

### Fos activation

All animals were sacrificed immediately after the last blood sample collection (120 min after commencement of FST) to allow assessment of expression of Fos in HPA axis- and stress-regulatory circuitry. Fos is the product of the immediate early gene c fos, and is reliably induced from largely undetectable levels by acute stressor exposure. We assessed the ability of the FST procedure to recruit neurons in the prefrontal cortex,(PFC) (prelimbic (PL) and infralimbic (IL) subdivisions), the lateral septum, ventral part (LSV), the paraventricular nucleus of the hypothalamus (PVN), and the subdivisions of the amygdala, central (CeAm), basolateral (BLA), and medial (MeAM).

We observed that CVS reduced Fos-immunoreactive (ir) nuclei in the males in the prelimbic cortex, STRESS F_(1, 26)_= 5.661, p<0.05, and in the infralimbic cortex, STRESS F_(1, 26)_=5.64, p<0.05 with no effect of drug treatment, nor interaction. (Figs. 4.B-C). Females did not show effects of adol CVS or C108297 in any PFC area (Table 1).

**Figure 4:**
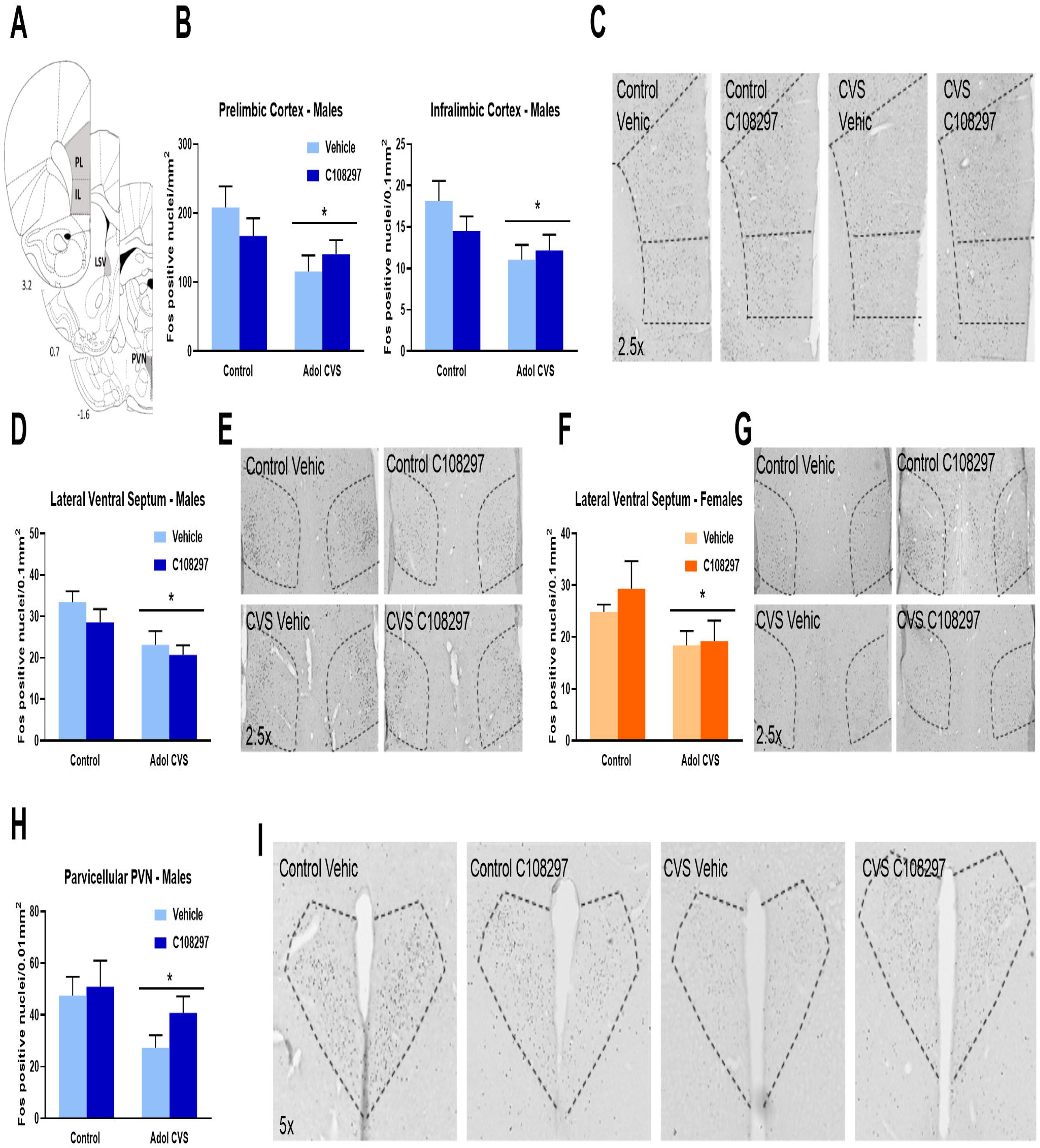
Fos immunoreactivity (Fos-ir) in the prelimbic and infralimbic divisions of the prefrontal cortex (PFC), lateral ventral septum (LSV) and parvicelullar division of the paraventricular nucleus of the hypothalamus (PVN). Animals were subjected to adolescent CVS (adol) and concomitantly administered with CORT108297 (30mg/Kg) or vehicle during 2 weeks starting at PND 46. Brains were collected 2h after the onset of FST, after 6 weeks of recovery from adol CVS. **(A)** Representative images from brain atlas showing the analyzed areas (Paxinos and Watson, 1998). **(B-C)** Fos-ir in the PFC of males and representative images. **(D-E)** Fos-ir in the LSV of males and representative images. **(F-G)** Fos-ir in the LSV of females and representative images. **(H-I)** Fos-ir in the PVN of males and representative images. Data are presented as mean ± s.e.m. *: significant result p < 0.05.

In the lateral ventral septum, adol CVS caused a reduction of Fos immunoreactivity in response to the acute stressor in both sexes: males, STRESS F_(1, 27)_=9.726, p<0.01; females, STRESS F_(1, 25)_=4.710, p<0.05, with no effect of C108297 or STRESS by TREATMENT interaction (Figs. 4.D-E and 4.F-G).

Fos-ir was reduced in the parvicellular region of the paraventricular nucleus of the hypothalamus in males (main effect of STRESS F_(1, 27)_=4.421, p<0.05) (Figs 4.H-I) but not in females (Table 1). There was no STRESS by TREATMENT interaction.

In the amygdala, there were main effects of STRESS in the medial amygdaloid nucleus (MeAm), F _(1, 27)_=10.36, p<0.01, the central amygdaloid nucleus (CeAm), F_(1, 27)_=6.846, p<0.05 and the anterior part of the basolateral amygdaloid nucleus (BLA), F_(1, 27)_=8.572, p<0.01. In the case of MeAM and BLA, planned comparisons revealed that the CVS-vehicle group had significantly less Fos immunoreactive nuclei than the control-vehicle group in these areas (p<0.05 respectively), while the CVS-C108297 group did not differ from the respective unstressed group. This suggests a role of GR signaling during adolescence in defining the long-term effect of CVS on amygdala excitability in males, particularly in the medial and basolateral subdivisions. Effects shown in (Figs. 5. B-C). There was no effect of stress or drug in females (Table 1).

**Figure 5:**
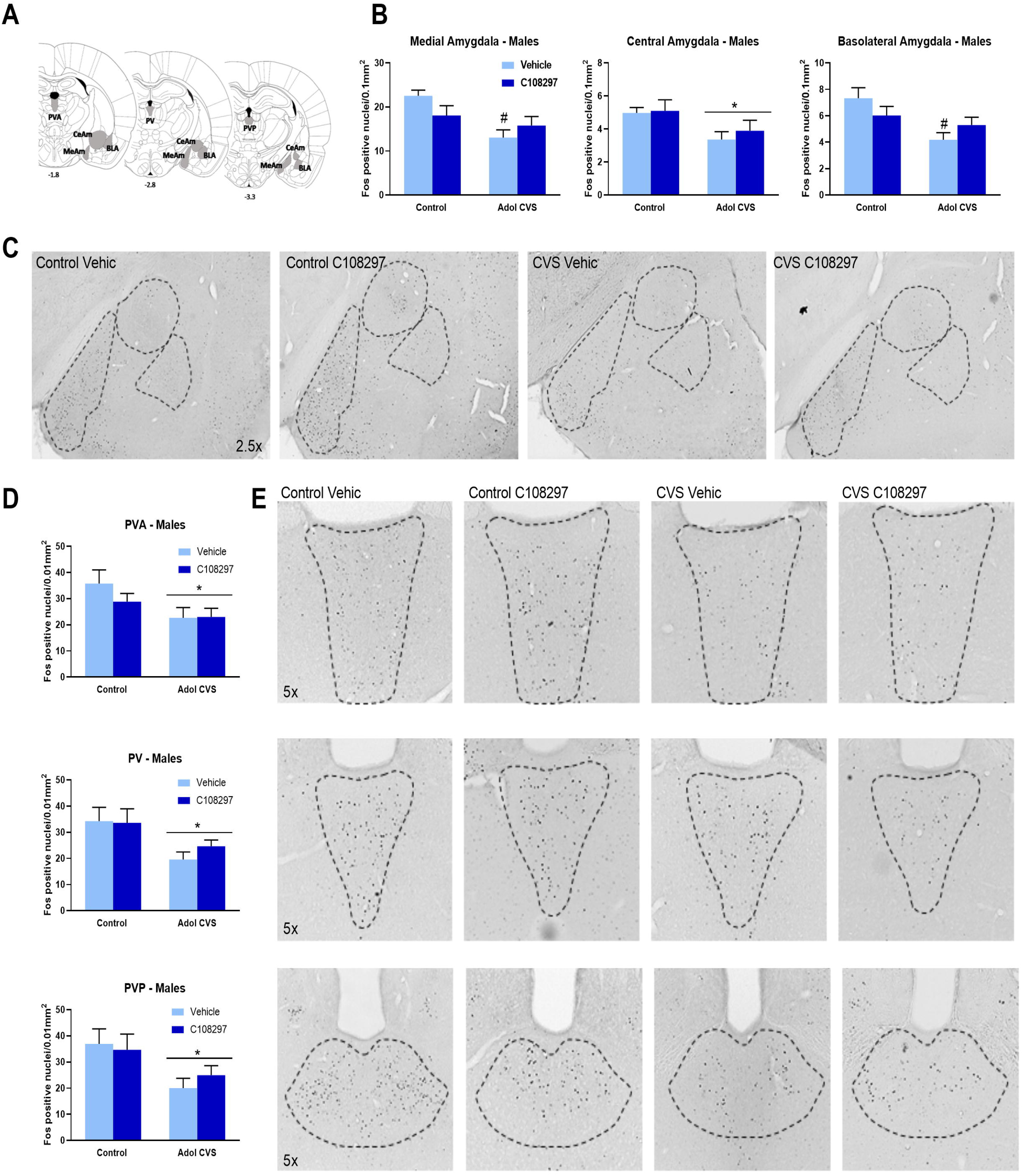
Fos immunoreactivity (Fos-ir) in the amygdala (medial (MeAm), central (CeAm) and basolateral divisions) and the paraventricular thalamic nucleus (anterior (PVA), intermediate (PV) posterior (PVP) divisions). Animals were subjected to adolescent CVS (adol) and concomitantly administered with CORT108297 (30mg/Kg) or vehicle during 2 weeks starting at PND 46. Brains were collected 2h after the onset of FST, after 6 weeks of recovery from adol CVS. **(A)** Representative images from brain atlas showing the analyzed areas (Paxinos and Watson, 1998). **(B-C):** Fos-ir in the amygdala of males and representative images. **(D-E)** Fos-ir in the paraventricular thalamic nucleus of males and representative images. Data are presented as mean ± s.e.m. *: significant result p < 0.05. ^#^: planned comparisons effect vs control vehicle group.

There was a reduction in number of Fos-ir nuclei in all the subdivision of the paraventricular nucleus of the thalamus: PVA, STRESS F_(1, 25)_=4.871, p<0.05; PV, STRESS F_(1, 27)_=8.162, p<0.01; PVP, STRESS F_(1, 23_ =7.812, p<0.05. (Figs. 5.D-E). No effects were observed in the females (Table 1), and there were no interaction effects in either sex.

## Discussion

The results of this study provide evidence supporting differential behavioral, physiological and stress circuit reactivity to adolescent chronic stress exposure in males vs. females, with pronounced sex differences evident in tests of emotional reactivity, coping behavior, HPA axis function and neuronal activation by stressors. In response to adolescent stress, males had heightened reactivity to novelty and exhibited marked reduction in neuronal excitation following swim stress in adulthood, whereas females developed a passive coping strategy and enhanced HPA axis stress reactivity. Only the latter effect was attenuated by treatment with the GR modulator C108297.

Glucocorticoids are known to have powerful effects on metabolism, and use of C108297 in adolescence had a marked impact on body weight gain during drug exposure as well as during the post-exposure period. Moreover, C108297 and adolescent stress produced an additive effective on body weight gain during the stress period. Since exposure to C108297 was systemic, it is possible that modulation of GR signaling during adolescence is able to program peripheral and/or central mechanisms regulating metabolism.

### Emotional reactivity in the open field test and coping in the forced swim test

Exposure to CVS in adolescence differentially affected behavioral responses in adult males and females. The avoidance of the center area and reduced exploration in an open field is typically interpreted as an indicator of heightened emotional reactivity to the novel environment (Faraji et al., 2014). These data are consistent with prior findings from our group and others (Cotella et al., 2019; Green et al., 2013; Ilin and Richter-Levin, 2009; Jacobson-Pick and Richter-Levin, 2010; Luo et al., 2014; McCormick et al., 2008; Zhang and Rosenkranz, 2012), and suggest that prior stress reorganizes behavioral responses to potential danger in males.

In the present study, performance in the open field test was not affected in females. These data contrast somewhat a prior study from our group (Smith et al., 2018). This discrepancy may relate to the CVS protocol used: in the present work, animals did not receive periodic overnight isolation or crowding, which may constitute a more profound stress exposure for females (Note that this was done to avoid stressing the animals outside the range of bioavailability of CORT108297). In addition, the daily injection of the GR modulator or vehicle during 2 weeks is another factor to take into account, given this constitutes itself a minor chronic stress situation that has been argued to evoke responses in the brain (Moghaddam, 2002).

In the case of the coping behavior during the forced swim test, we replicated previous results of increased immobility in adult females previously subjected to adolescent CVS (Wulsin et al., 2016) but in the case of the males, we only observed a modest effect of adolescent CVS at the end of the test, which was also consistent with previous results (Cotella et al., 2019) but in a less marked way. Bourke and Neigh indicated that females but not males are vulnerable to the effects of adolescent stress on the forced swim test as adults (Bourke and Neigh, 2011), and another study fails to report effects in male rats (Toth et al., 2008), suggesting that females may be more sensitive to the impact of adolescent stress on selection of coping strategy

It is notable that administration of C108297 did not block effects of CVS but did promote behavioral changes independent of stress. As was the case with body weight, it is possible that GR signaling during this developmental period has a role in defining coping strategies to aversive situations later in life.

### HPA axis regulation

Adolescent CVS reduced baseline AM ACTH in the males, but had no effect on corticosterone secretion. Though significant, the decrease in ACTH occurs in the low range of circulating ACTH, and may not be sufficient to significantly affect drive to the adrenal. In contrast, adolescent CVS increased ACTH and corticosterone in females, latter of which was blocked by treatment with C108297. Remarkably, C108297 reduced baseline levels of corticosterone in both sexes and prevented the increase in adrenal sensitivity caused by CVS.

The lack of effect of C108297 on ACTH results in both sexes suggests that the effects of drug on corticosterone levels might likely occurred by regulation of adrenal gland reactivity. This absence of effect on ACTH is consistent with prior findings indicating that C108297 does not regulate the expression of CRH in the PVN in intact rats (Zalachoras et al., 2013). Possible mechanisms of enhanced adrenal sensitivity may be primary hypertrophy of the gland or to increased splanchnic nerve activation, which regulates the reactivity of the adrenal cortex to ACTH (Engeland and Arnhold, 2005). Thus, modulation of GR signaling during adolescence causes lasting changes in HPA axis function, perhaps due to modification of adrenal development occurring in adolescent life. As discussed in (Russell D. Romeo, 2010a, 2010b), it is still unknown how adrenal function is developing during adolescence and how this is affected by stress. Nevertheless, based on human literature, we know that there is a general thickening of the adrenal cortex during this life epoch, which has been mostly studied in association to pubertal changes in sex hormones (Hui et al., 2009; Nakamura et al., 2015), while relevance for stress response remains unresolved.

Exposure to adolescent CVS does not affect the magnitude of the male HPA axis responses to an acute novel stressor in adulthood (5 weeks later) (Cotella et al., 2019; Smith et al., 2018). This contrasts with lasting sensitization seen in adults receiving CVS and tested 5 weeks later (Cotella et al., 2019), suggesting that if anything, adolescence confers resilience to the long-term effect of stress on the HPA axis in males. Unlike males, adolescent stress sensitized HPA axis reactivity in adult females, suggesting a lasting impact on stress reactivity. The data suggest adolescent stress may make females susceptible rather than resilient to subsequent stress. Of note, our prior study using a different CVS regimen (including social stressors) caused females to be hyporesponsive to acute stress, indicating differential reorganization of stress susceptibility (Wulsin et al., 2016). These data agree with those of Pohl et al. (Pohl et al., 2007) exploring two different adolescent chronic stress protocols of different intensity in female rats. These studies indicate that enhanced corticosterone secretion was observed in the group subjected to a subjectively “milder” stress protocol (no cold water stress). Collectively the data suggest that adolescent females exhibit long-term stress susceptibility across a broad range of stressor intensities, with the characteristics of stress regimen dictating the magnitude and valence of behavioral outcomes.

### C108297: Mechanism of Action

Administration of C108297 prevented HPA axis sensitization following CVS in females, indicating involvement of GR signaling. C108297 is known to have both agonistic and antagonistic properties in vivo (Viho et al., 2019; Zalachoras et al., 2013) and thus its valence of action is difficult to specific. Prior studies indicate that pre-stress exposure to C108297 reduces corticosterone secretion following forced swim in previously unstressed rats (Solomon et al., 2014), suggesting a GR agonistic action in this context. Moreover, Zalachoras et al note that C108297 treatment enhances passive avoidance performance memory in a manner similar to that of corticosterone suggesting an agonist effect (Zalachoras et al., 2013).

C108297 reduces body weight gain and absolute body weight in both sexes and this effect remained weeks after cessation of the treatment. Prior studies have reported body weight reduction following administration of C108297 and other GR targeting drugs (Asagami et al., 2011; Belanoff et al., 2010), indeed leading to trials testing efficacy in treatment of obesity. The prior studies attribute body weight effects to peripheral GR antagonist actions on metabolism. In our study, C108297 produces resistance to body weight gain during treatment as well as several weeks following discontinuation, consistent with an organizational action of adolescent glucocorticoids on metabolic trajectory.

### Mapping of neuronal recruitment in response to forced swim test

Quantifying Fos immunoreactive cells after exposure to FST provides insight into how adolescent stress can program recruitment of brain regions integrating the response to challenge. In general, adolescent stress blocked recruitment of several brain regions known to be involved in stress integration, including the prefontal cortex, the lateral ventral septum, the paraventricular nuclei of the thalamus and hypothalamus, and the central, medial and basolateral divisions of the amygdala. Consistent with our data, prior studies also note decreased prefrontal cortical activation using different modalities of adolescent stress and acute adult stimulus (Ishikawa et al., 2015). In adult males, fos expression to a novel environmental stressor is known to decrease in many of these brain regions immediately following a week of CVS, and this reduction is still observed in the prelimbic cortex and the lateral ventral septum after a month (Ostrander et al., 2010). Our results suggest that neurophysiological adaptations to prior stress experience may render stress-integrative circuitry less responsive to novel challenges. This stands in contrast to females, where (except for the lateral septum) adolescent stress is ineffective in modulating Fos reactivity following forced swim.

The combination of behavioral resilience and generalized reduced post-FST Fos expression in males suggests that adolescent stress may act to limit neural responses to subsequent stress and thereby buffer its overall impact. Prior studies indicate adolescent (male) HPA axis responses to both acute and chronic stress are more pronounced than adult responses (Jankord et al., 2011; Russell D. Romeo, 2010a; Viau et al., 2005), accompanied by higher Fos immunoreactivity in the PVN compared to adults (Viau et al., 2005). It has also been reported that prepubertal males have reduced Fos expression in the medial amygdala after an acute stressor compared to adults (Kellogg et al., 1998), indicating the relevance of developmental differences regarding the sensitivity of different brain regions in response to stress.

It is important to also consider the limited resilience evident in females. In our prior studies, the differential impact of adolescent stress (relative to adults) on the HPA axis is not observed in females (Wulsin et al., 2016). In combination with the generalized absence of adolescent stress reductions in Fos expression, the data suggest that reduced stress reactivity during adolescence in females (Russell D. Romeo, 2010a; Wulsin et al., 2016) (relative to males) may result in enhanced neurocircuit activation in adulthood. It is possible that the failure to robustly engage central stress pathways during adolescent stress is a factor in limiting stress resilience in females, resulting in more pronounced behavioral reactivity later in life.

### Conclusion

The results here presented support the idea that experiencing chronic stress during adolescence evokes sexual dimorphic endpoints. Furthermore, we also demonstrate that some of these enduring adolescent stress effects could be prevented by modulating the signaling through the glucocorticoid receptor during the exposure to stress, suggesting the involvement of these receptors in the adult phenotype observed. Based on the disparities in the effects observed between males and females, it is evident that either the organizational or the activational effects of sex hormones play a relevant part modulating the effects of stress during development and interacting with the glucocorticoid signaling. Interestingly, we also observed that modulating this receptor system in unstressed animals also affected some of the adult outcomes, evidencing the participation of glucocorticoid dynamics during this developmental phase as a programming factor for adult responses.

## Acknowledgements

We would like to thank Corcept Therapeutics for providing the Cort108297 compound. We also would like to acknowledge other members of Dr Herman’s laboratory and other researchers in the Reading Campus of the University of Cincinnati, especially Dr. Matia Solomon, for their support and helpful comments on the results.

