## Supplemental Figure 1 for "Selective modulation of the glucocorticoid receptor with CORT108297 during chronic adolescent stress evokes sex-specific effects in adulthood"

**Supplemental material**

**Total growth during the experiment:** In the case of the males, we observed main effects of TREATMENT F_(1,36)_=5.082, p<0.05, TIME_(9,324)_= 2718.27, p<0.001 that determined less absolute body weight in the animals treated with C108297 regardless of stress (p<0.05). There was also a TIMExTREATMENT interaction F_(9,324)_=9.18, p<0.001 and TIMExSTRESS interaction F_(9,324)_= 3.86, p<0.001.The analysis of planned comparisons showed that the control-C108297 group had continuous lower body weight than the control vehicle group starting on week 3 of experiment (p<0.05) and the CVS-C108297 group showed reduced body weight compared to the control vehicle group between weeks 2 and 5 and on weeks 7 and 9 (p<0.05). It is important to remark that there were no initial differences in body weight between groups so all the observed effects on absolute body weight are due to the experimental variables. Results shown in Figure S.1.A

In the females, there were main effects of TREATMENT F_(1,44)_=22.68, p<0.001 and TIME F_(9,396)_=1908.77, p<0.001 as well as an interaction TREATMENTxTIME F(_3,396)_= 29.27, p<0.001. The analysis of planned comparisons showed that the control group treated with C108297 had continuous reduced body mass since week 3 (p<0.05) while the group that was subjected to CVS and C108297 had the same effect starting on week 2 (p<0.05). There were no differences between these two groups (Fig S.1.B).

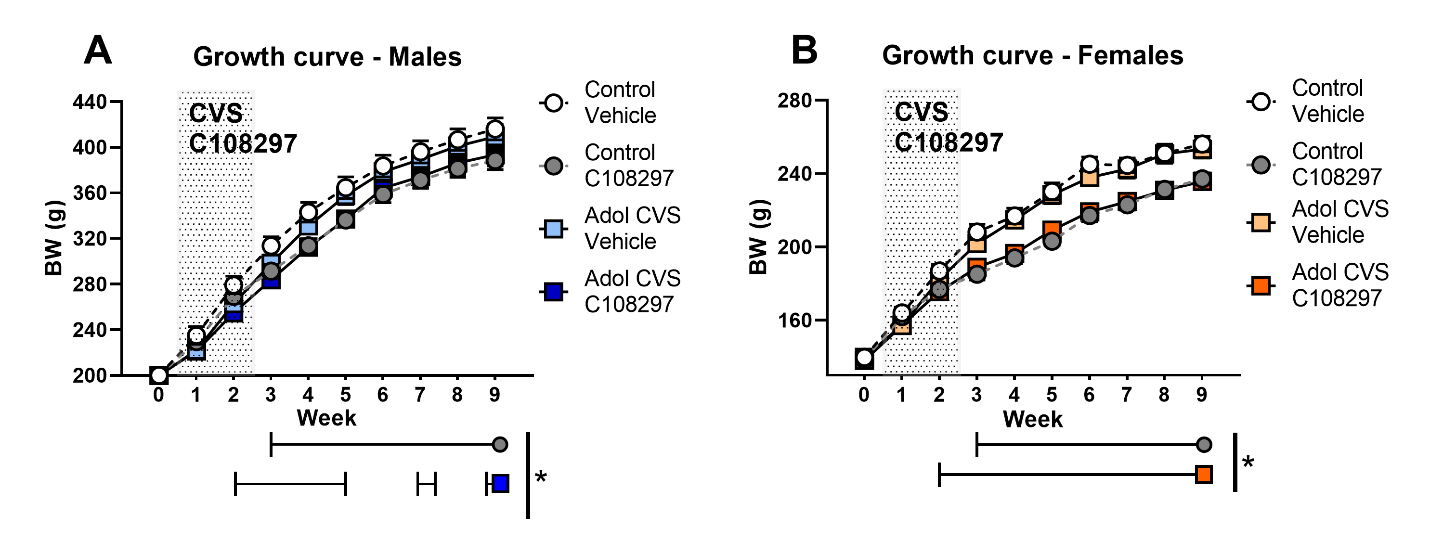
 **Figure S.1: Absolute weight across the experiment in males (A) and females (B).** Animals were subjected to adolescent CVS (adol) and concomitantly administered with CORT108297 (30mg/Kg) or vehicle during 2 weeks starting at PND 46. Body weight was taken weekly since before the beginning of CVS and daily during CVS. Data are presented as mean ± s.e.m. *: significant result p< 0.05 for planned comparisons versus Control-vehicle group.
