## Supplemental Table 1 for "Selective modulation of the glucocorticoid receptor with CORT108297 during chronic adolescent stress evokes sex-specific effects in adulthood"

| Variable | Group | Males | Females |
| --- | --- | --- | --- |
| Adrenal weight | Control - Vehicle  Control – C108297  CVS – Vehicle  CVS – C108297 | 35.46 ± 2.19  34.49 ± 2.07  37.88 ± 2.24  37.34 ± 2.08 | 41.59 ± 2.71  43.09 ± 2.65  43.53 ± 2.63  45.66 ± 2.69 |
| Thymus weight | Control - Vehicle  Control – C108297  CVS – Vehicle  CVS – C108297 | 465.88 ± 20.88  397.07 ± 19.81  426.59 ± 20.88  385.12 ± 19.81 | 305.72 ± 16.09  300.27 ± 15.20  278.24 ± 14.51  317.55 ± 14.57 |
| Heart Weight | Control - Vehicle  Control – C108297  CVS – Vehicle  CVS – C108297 | 1309.99 27.05  1273.88 28.63  1362.59 26.44 **]**  1347.03 26.43 **]** | 891.24 ± 14.42  888.11 ± 14.67  911.96 ± 14.01  896.57 ± 14.42 |

**Table S.1: Somatic markers of stress** Animals were subjected to adolescent CVS (adol) and concomitantly administered with CORT108297 (30mg/Kg) or vehicle during 2 weeks starting at PND 46. Organs were collected after euthanasia and wet weight was taken and covaried with bodyweight. Data are presented as mean ± s.e.m. ] represents main effect of CVS.
